# Inverse Game Theory Characterizes Frequency-Dependent Selection Driven by Karyotypic Diversity in Triple Negative Breast Cancer

**DOI:** 10.1101/2025.03.23.644809

**Authors:** Thomas Veith, Richard Beck, Joel S. Brown, Noemi Andor

## Abstract

Chromosomal instability (CIN), characterized by pervasive copy number alterations (CNAs), significantly contributes to cancer progression and therapeutic resistance. CNAs drive intratumoral genetic heterogeneity, creating distinct subpopulations whose interactions shape tumor evolution through frequency-dependent selection. Here, we introduce, ECO-K (Ecological-Karyotypes), an inverse game theory framework that infers subpopulation interactions from longitudinal single-cell whole genome sequencing data. Applying this approach to serially-passaged, triple-negative breast cancer (TNBC) cell lines and patient-derived xenografts (PDXs), we systematically identified frequency-dependent selection dynamics governed by karyotypic diversity. Notably, in one PDX lineage, we found consistent evidence of karyotypically defined subpopulations acting as interaction hubs, associated specifically with chromosome 1 loss and chromosome 14p gain. Our framework provides testable predictions of intratumoral ecological dynamics, highlighting opportunities to strategically target key subpopulations to disrupt tumor evolution.

## 1. Introduction

The evolutionary process that unfolds during tumor progression often renders failure in the clinic a *fait accompli* as natural selection promotes the expansion of therapeutically resistant subpopulations [1, 9]. Chromosomal instability, a hallmark of cancer associated with advanced disease and poor outcomes [12], is a powerful evolutionary process that rapidly generates genetic diversity. CIN results in ongoing copy number alterations which are pervasive across cancer types [13], affecting larger segments of the cancer genome than any other alteration [11]. While CNAs are known to alter a cancer cell’s fitness [13], response to treatment [16], and immunogenecity [13], it remains unclear how distinct karyotypes differ in phenotype or whether they engage in frequency-dependent interactions that influence tumor evolution.

Evolutionary game theory (EGT) models assume that each subpopulation’s growth rate is determined by interactions with other subpopulations, typically represented as a payoff matrix [6]. This approach views a tumor as an evolving ecosystem and studies how shifts in population composition influence cancer progression [7]. Examples of hallmarks of cancer studied with EGT include tumor-immune interactions [14], angiogenesis [2], invasion [4, 7, 5], microenvironment independence [8] and evasion of apoptosis [3]. In EGT models, frequency-dependence occurs when the fitness (payoff) of a cancer cells depends not only on its subtype but also on the subtypes of others.

Despite its utility, a major limitation of traditional EGT approaches is that they often rely on predefined interaction rules rather than inferring them directly from experimental data. Most insights from EGT in cancer revolve around how transformed cells interact with normal cells, whereas interactions between tumor subpopulations with distinct karyotypes remain poorly quantified.

To address this, we introduce an inverse game theory approach that infers payoff matrices directly from empirical data. By applying this method to karyotypically distinct subpopulations, we systematically evaluate frequency-dependent selection within tumors, revealing how removing specific subpopulations may enhance or impair overall tumor growth dynamics.

## 2. Materials and Methods

### 2.1. Defining subpopulations and replicate groups

git link: defining clones

#### 2.1.1. Copy Number Calling and Chromosome Arm Aggregation

We first processed single-cell copy number data to generate chromosome arm–level copy number profiles. For each input file (a segment-by-cell copy number matrix provided as a compressed CSV), the genomic bins were annotated with their corresponding chromosome and band information using a loci file. For each chromosome arm, the copy number was taken as the modal value across all segments mapping to that arm. The resulting arm-level copy number matrix was saved for subsequent clustering.

#### 2.1.2. Subpopulation Assignment via k-means Clustering

For each patient-derived xenograft or cell line, we aggregated the arm-level copy number matrices from all available single-cell datasets. Subpopulation (SP) identities were then determined by applying k-means clustering to the aggregated matrix. Specifically, for a given *k* (with *k* ∈ {2, .. ., 10}), clustering was performed (with 25 random starts), and the average Silhouette score computed. The optimal number of SPs *n* was selected as:

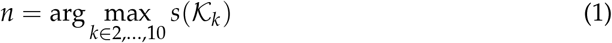

where *s*(𝒦_*k*_) is the average Silhouette score for the clustering with *k* centroids. Each cell is thereby assigned to a subpopulation (denoted *k*_*i*_, *i* = 1, …, *n*).

#### 2.1.3. Computing Subpopulation-Frequency Matrices and Replicate Grouping

Two serially-passaged lineages were considered members of the same biological replicate group if and only if they originate from the same PDX or cell line and have been exposed to cisplatin for a similar ratio of passages. For example, two lineages originating from the same PDX with different numbers of passages but similar relative fractions of passages exposed to cisplatin (e.g., 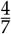 and 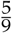 passages received treatment) would be considered replicates.

Using available metadata, each cell was annotated with its passage (timepoint) and an experimental label that reflects treatment exposure. Although SPs were defined over all cells from an origin, cells were further grouped by timepoint. For each timepoint *t* (or passage) within a replicate group, we computed a SP-frequency matrix 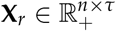, where each entry was defined as:

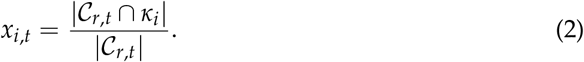

Here, 𝒞_*r*,*t*_ represents the set of cells in replicate *r* at timepoint *t*. Thus, *x*_*i*,*t*_ is the fraction of cells 𝒞_*r*,*t*_ that belong to the *i*^*th*^ subpopulation, 𝒦_*i*_.

### 2.2. Optimization Routine

To capture the dynamics of SP interactions and selection pressures, we modeled frequency-dependent selection using the replicator equation. This equation described the change in frequency of each SP over time, incorporating the fitness of each SP and the population-average fitness. For *n* SPs, the replicator equation is given by:

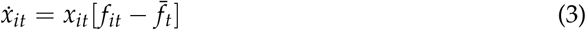

where *x*_*it*_ is the frequency of the *i*^th^ SP as defined in equation (2), *f*_*it*_ gives that SP’s fitness, and 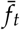 is the population-average fitness. The fitness of a given SP was defined as:

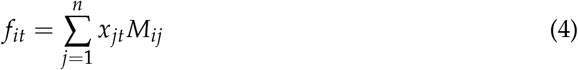

with payoff matrix entries (*M*_*i*,*j*_) where *m* is the payoff to an individual represented by row *i* when interacting with an individual represented by the column *j*.

Some interactions between SPs may have no fitness consequences. In that case the matrix entry representing their interaction would be set to zero. Non-zero interactions were inferred via the methods described in Section 2.4. The remaining interactions were estimated by minimizing the negative log-likelihood with the MatLab function fmincon using the sequential quadratic programming (SQP) algorithm within the MultiStart function framework.

The optimization routine was performed for multiple replicates simultaneously. Let Ω := {**X**_**r1**_, **X**_**r2**_, .. . } be the set of replicates within a replicate group, with entries as defined in equation (2). A subpopulation which never exceeds 10% frequency ∀*t* ∈ *T* in any replicate was excluded from downstream analysis. Suppose one such SP exists, then *n* := *n* − 1. Note that this leads to the payoff matrix *M* having identical dimensions across all replicates of a given group during each iteration of the optimization routine.

The probability of observing replicate **X** ∈ Ω given the payoff matrix *M* can be modeled by a normal distribution of residuals:

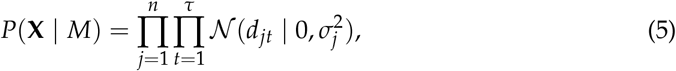

where *d*_*jt*_ = **X**_**jt**_ - *x*_*jt*_ represents the residuals, with **X**_**jt**_ being the observed frequency of SP *j* at timepoint *t*, and *x*_*jt*_ being the predicted frequency based on solution to Eqn. 3 given the payoff matrix *M*, and initial conditions 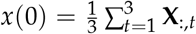. The variance 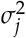 for each SP *j* was estimated as the standard deviation of the residuals across all time points:

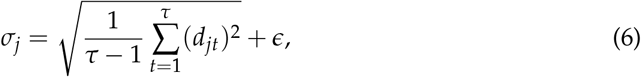

where *ϵ* is a small constant added to avoid division by zero. The overall log-likelihood for replicate **X** is then:

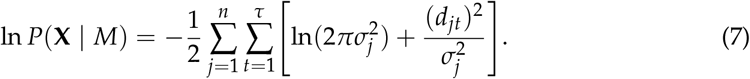

The cumulative negative log-likelihood across all replicates was calculated as:

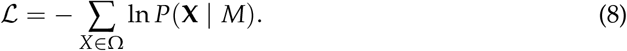

git link: [optimizer](https://github.com/MathOnco/EVO_K/blob/main/ecological_karyotypes/ecological_karyotypes.m)

### 2.3. Parsimonious Parametrization Based on Iterative BIC Calculation

We used the above optimization routine to select between models of different complexity as follows. We calculated the Bayesian Information Criterion (BIC) and corrected Akaike Information Criterion (AICc) across all replicates within a given group. These were given by:

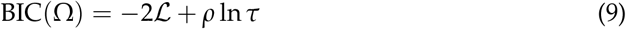

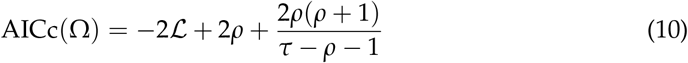

where ℒ is the negative log-likelihood calculated in section 2.2, *ρ* is the number of interactions in the model and *τ* is the total number of passages across all replicates.

During each iteration, the current set of non-zero interactions was estimated (see section 2.2), and the corresponding BIC and AICc calculated. The function then attempts to improve the model by considering setting each non-zero matrix entry to zero one at a time. For each matrix entry considered, the optimization process was repeated, and the BIC/AICc was recalculated. If setting a matrix entry to zero results in a lower BIC, indicating a better model fit with fewer interactions, the matrix entry was permanently set to zero. Even if setting the matrix entry to zero alone does not lower the BIC, we further test setting it to zero in combination with the removal of other matrix entries. This iterative removal continues until no further reduction in BIC was possible.

git link: [Likelihood function](https://github.com/MathOnco/EVO_K/blob/main/ecological_karyotypes/likelihood_function.m)

### 2.4. Correlation Between Growth Rate and Subpopulation Frequency

Recall that *T* = {*t*_1_, *t*_2_, …*t*_*t*_} is defined as all timepoints (i.e., passages) across replicates for a given replicate group Ω := {**X**_**r1**_, **X**_**r2**_, .. . }.Then, ∀*t* ∈ *T*, we defined the *growth rate α*_*i*,*t*_ of SP *i* between passages *t* and *t* + 1 by the difference in the natural logarithm of its frequencies, normalized by the difference in days:

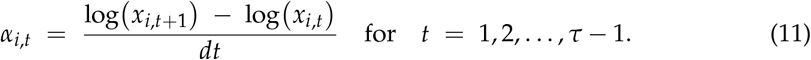

, where *d*_*t*_ is the difference between passage *t* and the subsequent passage in units of days. For each pair of subpopulations (*q, v*), we sought to determine whether the frequency of SP *v* influenced the growth rate of SP *q*. We therefore computed the Pearson correlation coefficient *R*_*qv*_ by correlating:

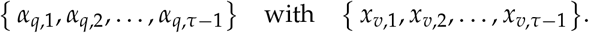

Hence,

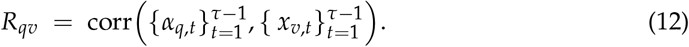

We used a *t*-test to obtain a corresponding *p*-value *P*_*qv*_ for the null hypothesis of zero correlation.

We then applied a Bonferroni correction across all *n* × *n* pairwise tests, i.e.,

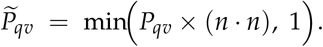

We define a *significance score* 𝒮_*qv*_ by combining the (Bonferroni-corrected) *p*-value 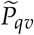 and the magnitude of the correlation:

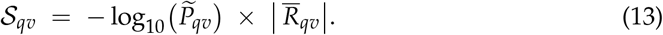

Higher scores indicate stronger frequency-dependent interaction between SPs *q* and *v*.

Following [10], we impose an upper limit on the number of potentially non-zero interactions. Specifically, if there were *τ* passages in the dataset, at most *τ* − 1 such pairwise interactions were retained as candidates for being non-zero in the payoff matrix. To implement this threshold, we rank the (*q, v*) pairs in descending order of 𝒮_*qv*_ and select the top *τ* − 1. All other off-diagonal entries in the payoff matrix were then set to zero. The sign of the correlation (±) was used to determine whether the selected payoff entry should be positive or negative, ensuring that both the magnitude and direction of the interaction were captured.

git link: test for frequency-dependent effects in growth data

### 2.5. Bootstrapping

To ensure the reliability of matrix entry estimates, we employed a subsampling-based method. For each **X** ∈ Ω, we generated *β* subsampled data sets 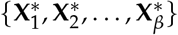 by randomly selecting unique time points (i.e., sampling without replacement) from the solution to Eqn. (3) using the payoff matrix **M** found via ECO-K. Specifically, each subsample 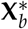 was formed by drawing a random subset of time points and the corresponding solution trajectories from the replicator equation.

For each subsampled data set 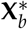, we performed the optimization routine as described in Section 2.2, which resulted in subsampled payoff matrices 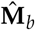 for *b* = (1, 2, .. ., *β*). After processing all subsampled data sets, we analyzed the resulting payoff matrix estimates 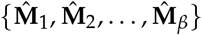. We calculated the means, standard errors, p-values, and confidence intervals of these estimates to assess their variability and statistical significance.

In particular, the mean and standard error of the estimated payoff matrix entries were given by:

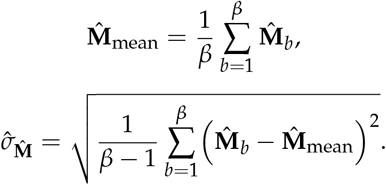

The 95% confidence intervals can be calculated using the percentiles of the subsampled distribution or using the standard error:

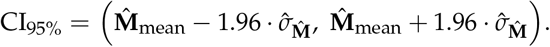

We tested the null hypothesis that a matrix entry was effectively zero by examining its p-value, calculated as a t-test comparing the distribution of 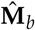 to zero.

git link: [bootstrapping function script](https://github.com/MathOnco/EVO_K/blob/main/bootstrapping/bootstrap_func.m)

### 2.6. Synthetic dataset generation

To test the robustness of our method, we generated 1000 artificial datasets with varying numbers of SPs and noise levels. For each dataset, we first randomly selected the number of SPs (*n*, where 2 ≤ *n ≤* 5) and assigned initial SP frequencies **X**_**0**_ = {**X**_**i**,**0**_, **X**_**j**,**0**_, .. ., **X**_**n**,**0**_**}** drawn from a uniform distribution, normalized such that they sum to 1.

Next, we generated a random payoff matrix *M* by drawing entries uniformly from the interval [−1, 1]. To restrict interaction complexity while still allowing for a sufficient number of nonzero interactions, we kept the top 2*n* − 1 (capped at 10) matrix entries by absolute value and set the rest to zero. We then solved the replicator dynamics (Eqn. 3) from *t* = 0 to *t* = 50 with these initial conditions. To ensure that the terminal state was at or near an Evolutionarily Stable Strategy (ESS), we required:

- that the difference between the maximum and minimum fitness values (rounded to one decimal place) at the end of the simulation be at most 0.1:

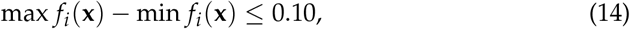
- and that all non-trivial eigenvalues of the Jacobian matrix,

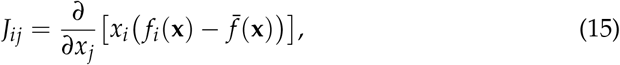

had negative real parts (as checked numerically).

If these criteria were satisfied, we regarded the final state of the replicator dynamics as nearing convergence to an ESS. While each system converged near an ESS, the speed at which it converged varied across artificial datasets (Supplementary Fig. A1).

From the simulation results, we focused on timepoints up to *t* = 30 and sampled 22 equally spaced points over this interval. At each of these timepoints, we recorded the SP frequencies **x**_**jt**_ ∈ **X**. We then introduced observational noise *ω* drawn uniformly from [0, 0.2] (i.e., *ω* ∈ [0, 0.2]), and perturbed each entry **x**_**jt**_ with an i.i.d. Gaussian random variable *η*_*jt*_ ∼ 𝒩 (0, *ω*). Specifically, the “noisy” version **X′** was computed element-wise as:

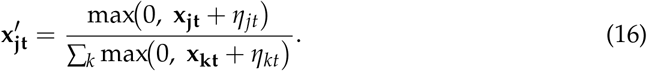

Thus, any negative values after noise addition were set to zero, and each timepoint was subsequently renormalized so that SP fractions sum to one. Finally, we applied our payoff recovery procedure to infer a payoff matrix *M***′** from the noisy dataset **X′**.

To assess how accurately we could infer the original payoff matrix, we used the following two performance metrics:

- 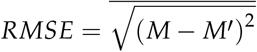, the usual root mean square error including all payoff matrix entries (Fig. 2), or just the non-zero entries (Supplementary Fig. A2).
- 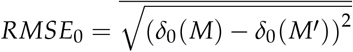, where *δ*_0_(·) is an indicator that is 1 if the corresponding entry of the matrix is zero (and 0 otherwise). This measures how well the zero vs. non-zero structure of *M* was recovered (Supplementary Fig. A3).

As a reference, the expected *RMSE* between two matrices whose entries are drawn independently from [−1, 1] can be approximated by considering the variance of their difference (under a uniform assumption), which is 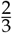. Hence, the expected *RMSE* between the randomly generated matrix and the one inferred via ECO-K was 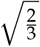.

### 2.7. Longitudinal single-cell whole genome sequencing data

Recently, researchers serially-passaged the triple-negative breast cancer cell line 184-hTERT to characterize subpopulation dynamics [17]. They employed CRISPR-CAS9 to derive a p53 knock-out (KO) version of the 184-hTERT cell line from the parental p53 wild-type (WT) cell line. The parental (WT) line was cultured for 60 passages and sampled four times to infer copy number status via single-cell whole genome sequencing (1599 cells sequenced on average, ranging from 1135-2061, 6395 total). Similarly, the KO cell line was cultured for 60 passages, with a replicate KO dish established at passage 20, which resulted in two KO lineages which were sampled for WGS a total of six times, establishing single-cell genomes for 1200 cells per passage (range of 728-1738 cells, 16795 cells total). None of the *in-vitro* experiments involved drug exposure.

Expanding their study to *in-vivo* models, the researchers established one HER2+ and three TNBC patient-derived xenograft (PDX) tumor mouse models. From these four PDX tumors, a total of 27 lineages were established. Among these, six serially-passaged PDX tumors were never exposed to drugs, while 18 were exposed to cisplatin in at least one passage. The PDX tumors were passaged between four and seven times over a range of 353 to 927 days, with single-cell genomes established at each passage for an average of 1146 cells originating from TNBC PDX mouse models (range of 466-1845 cells, 95954 total), and 890 cells from the HER2+ PDX (ranging from 497-1831 for a total of 8009 cells).

## 3. Results

### 3.1. Inverse game theory infers subpopulation interactions from interference patterns

We developed a comprehensive framework to detect and analyze frequency-dependent interactions between SPs in genomically resolved time-series data (Figure 1). Briefly, we first flagged SP pairs as interacting by testing whether the growth rate of one SP correlated with the frequency of the other SP [15]. We then optimized a payoff matrix (where each entry reflects the interaction strength for the corresponding SP pair) by minimizing a negative log-likelihood function under a multi-start strategy (Eqn. (7)). After this optimization, we iteratively removed individual entries from the payoff matrix if their removal lowered the Bayesian Information Criterion (BIC), thereby pruning insignificant matrix entries (Methods 2.3). Finally, we performed a bootstrap procedure on the reduced payoff matrix to determine which of the remaining interaction coefficients were significantly different from zero (Methods 2.5). This process produced a final matrix of frequency-dependent effects that was both parsimonious and supported by the data.

**Figure 1.**
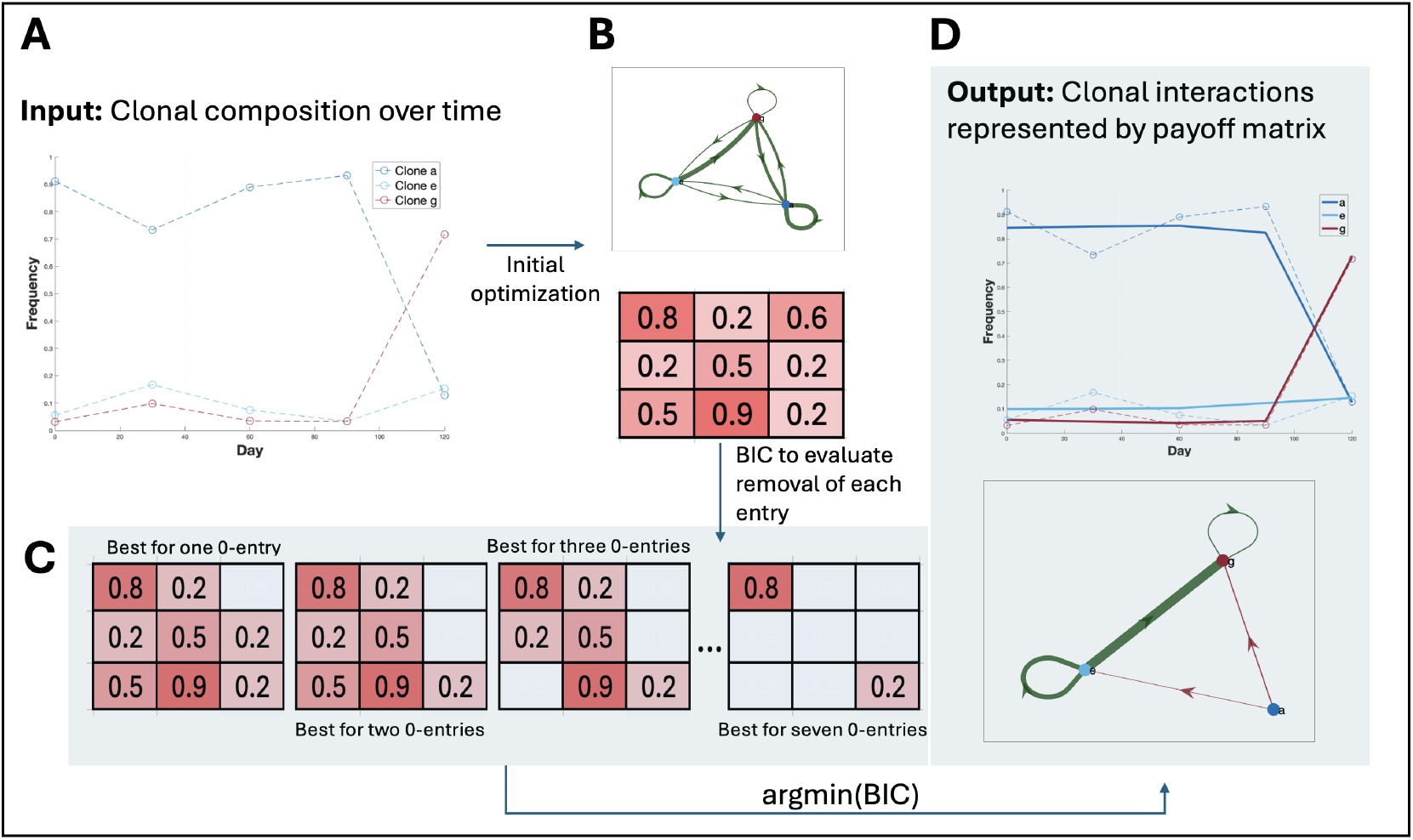
Overview of the optimization routine used in our method. The flowchart depicts the step-by-step process of optimizing the payoff matrix for SP interactions using ECO-K. The routine begins by initializing the interactions and setting up the optimization problem. An initial optimization was performed, and the Bayesian Information Criterion (BIC) was calculated with all interactions. The method then evaluates the removal of matrix entries to determine if this improves the BIC score. All indices in the matrix were tested for removal, but only the one that gives the lowest BIC score was retained. This routine was designed to ensure the most parsimonious set of interactions was used to capture the subpopulation dynamics.

To evaluate performance of the framework, we used a random payoff matrix *M*^*^, with entries 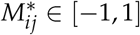. Using *M*^*^ we generate an artificial time series dataset Ω^*^(*ϵ, n*), with variable total number of subpopulations *n* and noise levels *ϵ* (Methods Section 2.6). For each artificial dataset, we compared the payoff matrix inferred by our framework *M*, to the underlying matrix *M*^*^ (Fig. 2). The results from these simulations indicate that our method is robust across a broad range of conditions. Even as the number of SPs and noise levels vary, and despite the random initialization of frequencies, the recovery algorithm consistently infers the underlying payoff structure better than would be expected by random chance (Methods 2.6).

**Figure 2.**
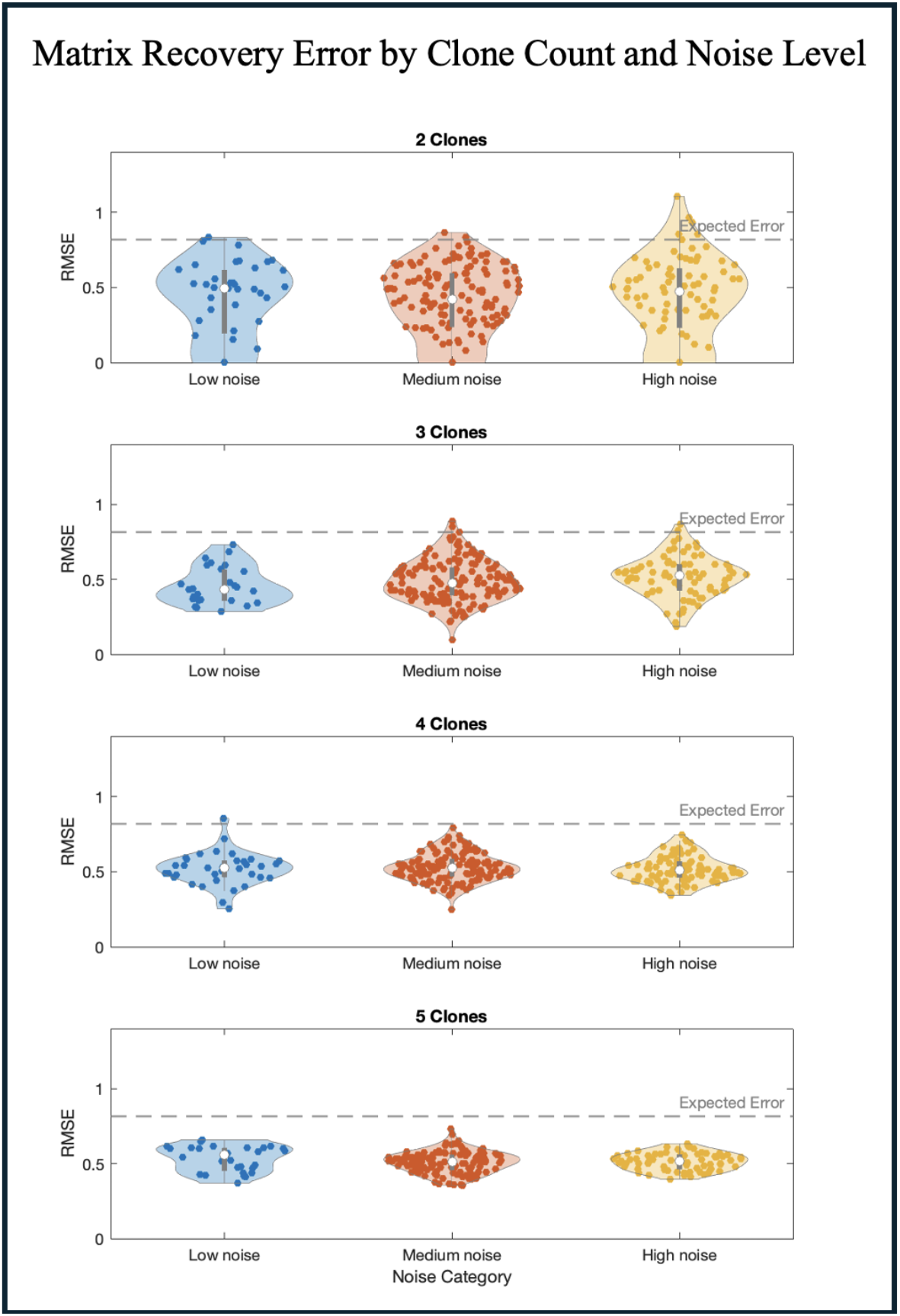
Payoff matrix recovery error as a function of noise and subpopulation count. Violin plots illustrating the distribution of recovery errors across 1000 artificial datasets, grouped by both the number of SPs present in the simulation (2 to 5 SPs) and the level of noise introduced. Noise levels were categorized as low (0.00–0.03), medium (0.03–0.13), or high (0.13–0.20).

### 3.2. Evaluating evidence for frequency dependent selection in TNBC

Copy number alterations play a critical role in cancer development and progression, impacting cellular phenotypes and patient outcomes. Shah and colleagues [17] recently performed serial-passaging experiments followed by single-cell DNA sequencing of triple-negative breast cancer cell lines and patient-derived xenografts (PDXs) (Table 4), allowing for longitudinal copy number inference. *In-vitro* and *in-vivo* lineages were cultured for 20-60 passages and sampled four to seven times for sequencing and copy number inference. A subset of lineages were exposed to cisplatin at various passages to evaluate whether the drug selects for certain copy number alterations.

We used ECO-K to evaluate evidence for frequency-dependent selection in these cell lines and PDXs. We grouped 27 lineages into 13 replicate groups (Table 4). Two lineages **X**_**i**_ and **X**_**j**_ were considered replicates (i.e. members of the same replicate group), if and only if they originate from the same PDX or cell line and have been exposed to cisplatin for a similar ratio of passages (Section 2.1). We clustered single-cell genomes across all replicates within a given group by their karyotype to define SPs (Section 2.1), which resulted in an average of four SPs (between two and seven) per group (Fig. 5). Thus if we detected interactions between every pairwise set of SPs, ECO-K would find 208 non-zero entries across 13 payoff matrices.

Applying ECO-K to these 13 replicate groups, we identified 129 interactions in total across all groups, with an average of five interactions per group (one group had no interactions, the most interactions detected in a replicate group was eight) (Table 4). Bootstrapping confirmed that 120 (93%) of these interactions exhibited a consistent directional trend, being predominantly positive or negative (Table 4). We considered a replicate group to have evidence of frequency-dependent selection if at least one replicate had an error below 0.05 and all replicates had an effect size above 0.1. Out of 13 replicate groups, six met these criteria.

To clarify whether intrinsic growth differences alone between subpopulations (SPs) could explain our observations, we compared the ECO-K framework introduced here to the FitClone method [17]. FitClone uses a Bayesian Wright-Fisher diffusion model, which estimates constant growth rates (referred to as selection coefficients) for each SP independently. Importantly, FitClone assumes that the growth rate of each SP is unaffected by the presence or frequency of other SPs within the tumor. In contrast, ECO-K explicitly captures frequency-dependent selection through a payoff matrix, where each entry quantifies how the presence of one SP affects the growth of another. Non-zero entries in this payoff matrix represent meaningful interactions between specific pairs of SPs, while zero entries imply no interaction. Here, each SP represents a distinct karyotypic configuration, analogous to a “strategy” in evolutionary game theory.

Among six replicate groups with evidence for frequency dependent selection, two could be better explained by the FitClone model, and two were better explained by ECO-K, while two had insufficient SPs (must be >2) to be evaluated by FitClone (Table 4). Here we highlight one lineage with strong evidence of frequency-dependent selection (Figure 5): Replicate Group 4 in PDX TNBC-SA1035. The most notable changes occur between days 150 and 250 (Fig. 5A), corresponding to the trajectory where the dominant subpopulation transitions from SP A to SP C. This is illustrated in Fig. 5B (bottom panel) as the system transitions from the forward-facing plane to the right-facing plane.

In Replicate Group 4, the best fit payoff matrix for the four SPs had six non-zero entries. Potential mechanisms for interactions arise from analyzing the replicator dynamics and copy number profiles of the subpopulations. Subpopulation F was the only subpopulation that interacted with all three other SPs and was characterized by chromosome 1 loss and gain of chromosome 14p (Fig 5 B-D). Chromosome 1 harbors multiple genes involved in metabolism (e.g., GLUL, involved in glutamine metabolism and NADPH-dependent enzymes). Its loss might impair SP F’s ability to synthesize certain metabolites, making it more dependent on metabolic exchange with other SPs, facilitating broad interactions.

Alternatively, if we consider specifically SP F’s interaction with SP C: Subpopulation F’s gain of 14p may promote SP C due to upregulation of TGFB3, a signaling factor that supports proliferation and immune evasion. This compensates for SP C’s loss of 8q, which may have impaired its autonomous survival metabolic flexibility.

Collectively, these results support the hypothesis that karyotypic alterations can drive frequency-dependent interactions in specific contexts (Fig. 4).

**Figure 3.**
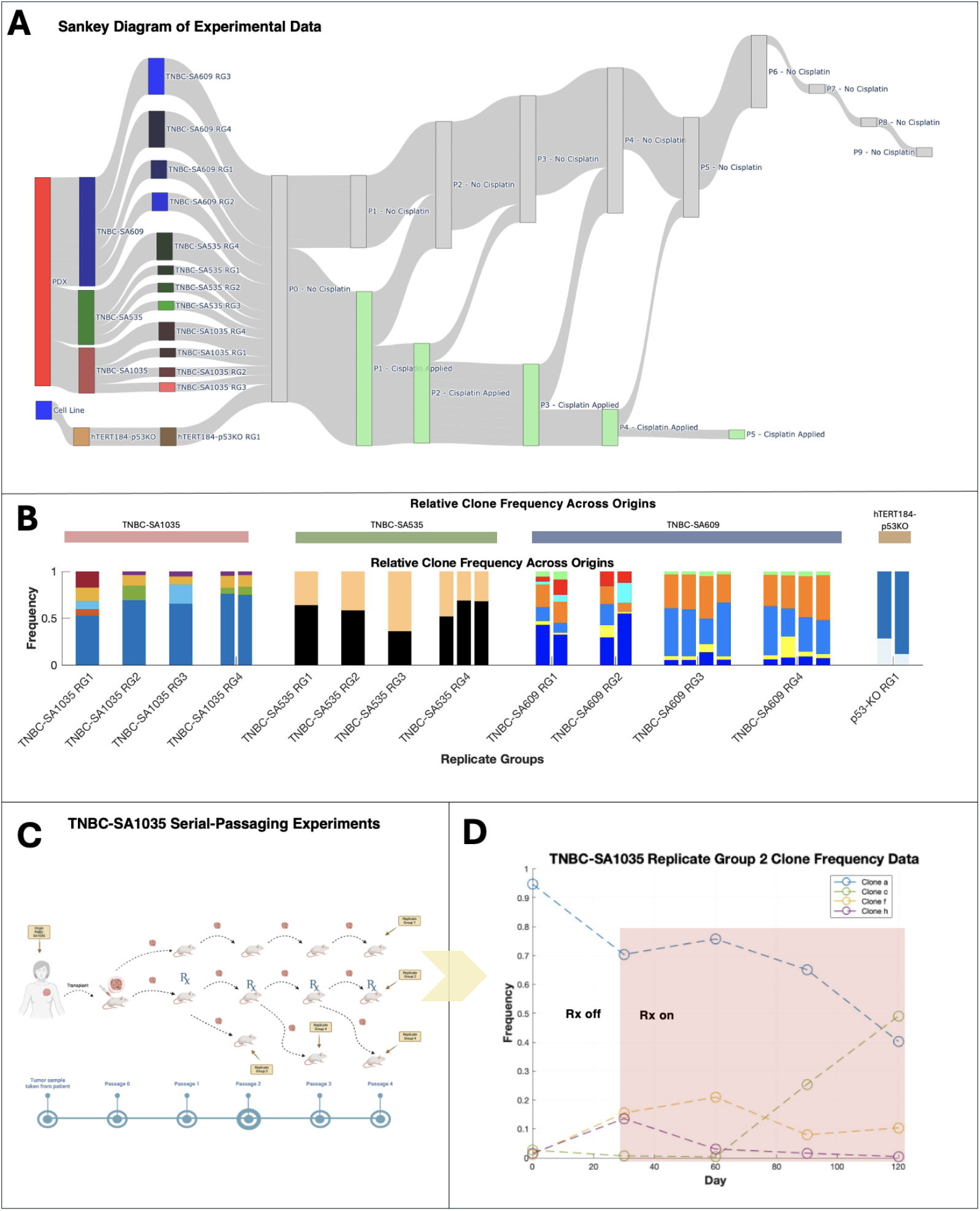
Overview of the serial-passaging experiments and example subpopulation dynamics in the SA1035-TNBC treatment arm. (**A**) Sankey diagram showing data origin (cell line or PDX), replicate groups, and cisplatin treatment schedules. (**B**) Relative subpopulation (SP) frequencies across origins and replicate groups, with colors indicating SP identities unique to each replicate group. (**C**) Schematic of serial-passaging in the TNBC-SA1035 patient-derived xenograft (PDX), illustrating tumor passaging under varying cisplatin regimens (treatment, no treatment, or holiday).(**D**) Example SP dynamics from TNBC-SA1035 Replicate Group 2, highlighting SP frequency shifts over time. Shaded regions indicate cisplatin administration.

**Figure 4.**
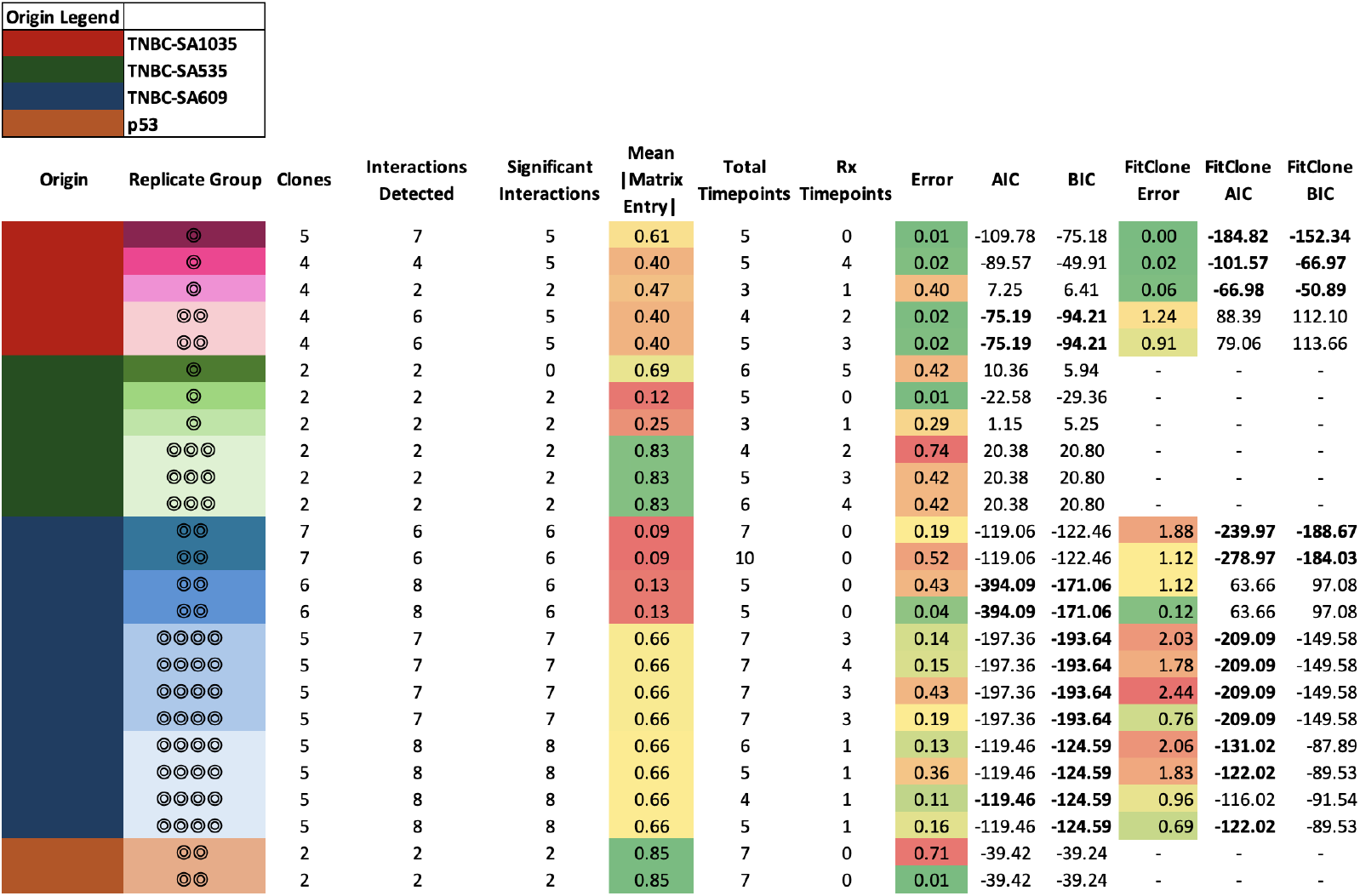
Comparison of ECO-K vs FitClone performance across various TNBC lineages. The table summarizes error, corrected Akaike Information Criterion (AICc), and Bayesian Information Criterion (BIC) for both ECO-K and FitClone models. Each row represents a replicate group, with color indicating sample origin. NA values indicate cases where model fits were not applicable or insufficiently supported. Overall, ECO-K provided better fits (lower AICc and BIC) in 5 cases, FitClone was superior in 5 cases, and the remaining 7 cases showed mixed results.

## 4. Discussion

In this study, we introduced a novel inverse game theory framework capable of inferring frequency-dependent interactions among karyotypically distinct tumor subpopulations. This approach leveraged single-cell whole genome sequencing data, which enabled the identification of key interactions that may drive a tumor’s evolutionary trajectory. Our results indicate that certain subpopulations may act as interaction hubs, driving ecological dependencies within the tumor microenvironment (Fig 5 B-C). Such insights underline how selective targeting of these hubs could strategically disrupt tumor progression.

**Figure 5.**
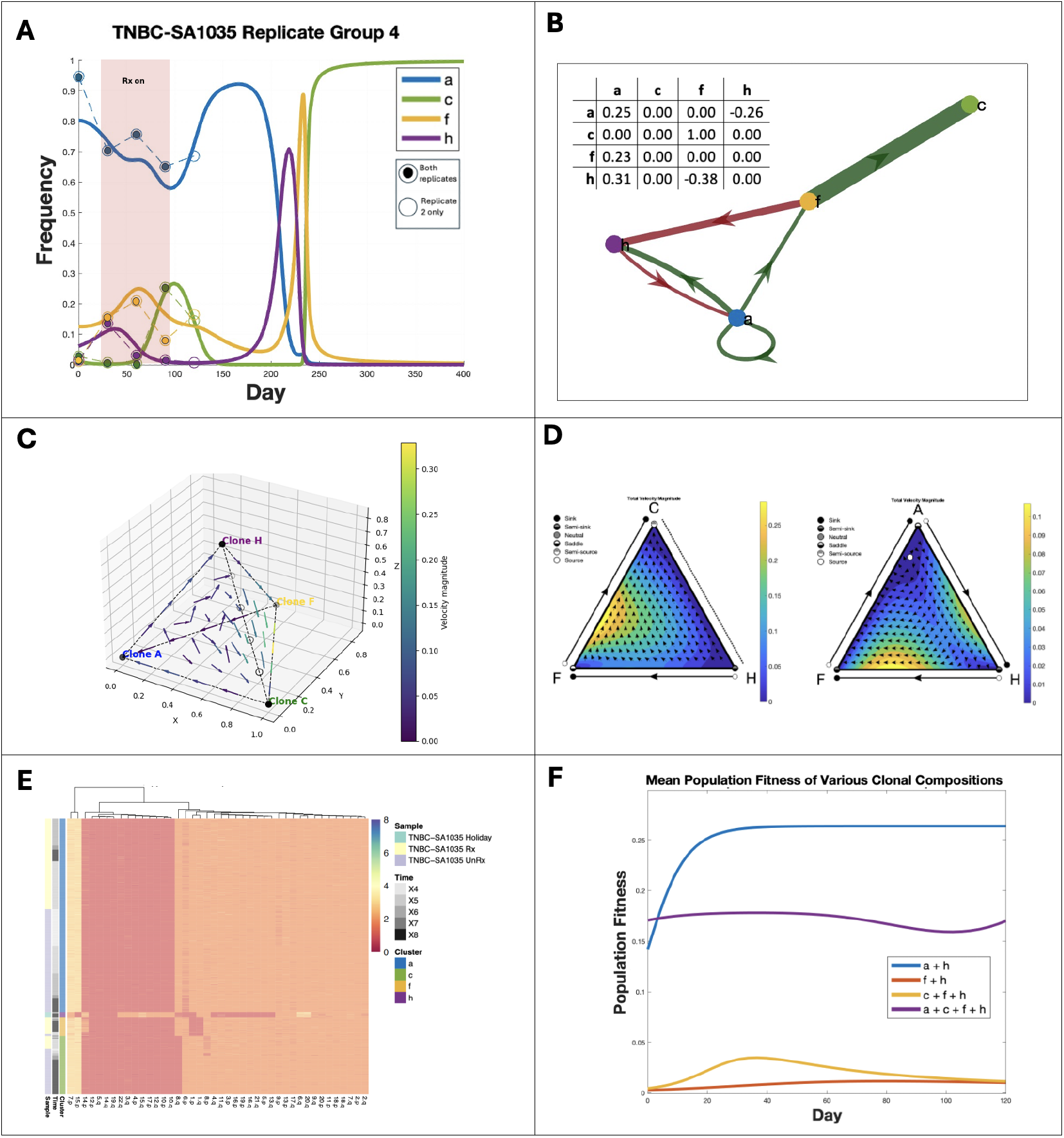
Frequency dependent interactions between karyotype defined subpopulations in TNBC PDX mouse models. (**A**) Observed (dashed lines) and replicator equation–predicted frequency dynamics (solid lines) for subpopulations (SPs) in TNBC-SA1035 Replicate Group 4, showing convergence to an evolutionarily stable state (ESS). (**B**) Payoff matrix diagram illustrating interactions between SPs. (**C**) Replicator phase diagram correlating to (B) (**D**) Ternary plots of replicator dynamics for selected SP combinations: C–F–H (left) and A–H–F (right). (**E**) Heatmap of copy number alterations (CNAs) across SPs, grouped by sample, time, and cluster. (**F**)Mean population fitness trajectories under different SP compositions. Removing specific SPs alters overall fitness relative to the original composition (purple).

However, our study does carry limitations which should be considered. Firstly, the inferred interactions and payoff matrices rely heavily on available longitudinal datasets, which often have limited temporal resolution and relatively few time points. Such data sparsity can constrain the robustness of inferred interactions and reduce the ability to detect subtle or transient frequency-dependent effects. Moreover, experimental variability between replicate groups and potential technical noise in sequencing data could further obscure genuine biological interactions or falsely imply significance.

Addressing the limitations would require experiments specifically designed to validate frequency-dependent interactions driven by karyotypic alterations. Future studies could adopt targeted perturbation experiments, involving the removal of specific subpopulations to observe impacts on tumor growth rates and evolutionary trajectories. This could be achieved through the administration of cisplatin, which has differential efficacy across karyotypes [17], that would provide the opportunity to evaluate the existence of SPs acting as interaction hubs (Fig. 5 B-C) and validate our predictions regarding changing tumor growth dynamics as a function of subpopulation composition (Fig. 5 E). Enhanced experimental resolution through higher temporal sampling and integration of multimodal data (e.g., transcriptomics, metabolomics) would also deepen our understanding of frequency dependent interaction dynamics and the phenotypic consequences of karyotypic alterations in tumor evolution.

Our inverse game theory pipeline suggests that frequency-dependent selection between cancer subpopulations, potentially influenced by distinct karyotypic profiles, may be empirically assessed *in-vivo*. The inferred fitness values could help guide therapeutic strategies aimed at exploiting or mitigating intratumor ecological dynamics by strategically targeting key subpopulations to hinder cancer progression.

## Author Contributions

Conceptualization, T.V. and N.A.; methodology, T.V., R.B. and N.A.; software, T.V. and N.A.; formal analysis, T.V. and N.A.; investigation, T.V. and N.A.; resources, N.A.; data curation, T.V., R.B.,. and N.A.; writing—original draft preparation, T.V. and N.A.; writing—review and editing, T.V. and N.A.; visualization, T.V. and N.A.; supervision, N.A.; project administration, N.A.; funding acquisition, N.A.

## Funding

This work was supported by the NCI grants 1R03CA259873-01A1, 1R37CA266727-01A1 and 1R21CA269415-01A1 awarded to N. Andor. The funders had no role in study design, data collection and analysis, decision to publish, or preparation of the manuscript.

## Acknowledgments

We thank Sohrab Salehi and Sohrab Shah for generously sharing the single-cell whole genome sequencing data from their work [17] and for providing insightful guidance on the details of preprocessing the data.

## Institutional Review Board Statement

Not applicable.

## Informed Consent Statement

Not applicable.

## Data Availability Statement

All code and data inputs are available at https://github.com/MathOnco/ECO_K.

## Conflicts of Interest

The authors declare no conflicts of interest. The funders had no role in the design of the study; in the collection, analyses, or interpretation of data; in the writing of the manuscript; or in the decision to publish the results.

## Abbreviations

The following abbreviations are used in this manuscript:

CIN: chromosomal instability
CNA: copy number alteration
EGT: evolutionary game theory
IGT: inverse game theory
PDX: patient derived xenograft
scWGS: single cell whole genome sequencing
TNBC: triple negative breast cancer

## Disclaimer/Publisher’s Note

The statements, opinions and data contained in all publications are solely those of the individual author(s) and contributor(s) and not of MDPI and/or the editor(s). MDPI and/or the editor(s) disclaim responsibility for any injury to people or property resulting from any ideas, methods, instructions or products referred to in the content.

## Notes

https://github.com/MathOnco/ECO_K

### Competing Interest Statement

The authors have declared no competing interest.

### Summary of Updates

- github repository links of associated code

## References

[1] PC Nowell. “The clonal evolution of tumor cell populations”. In: Science (New York, N.Y.) 194.4260 (Oct. 1976). 00000, pp. 23–28. ISSN: 0036-8075.

[2] J. Folkman. “The role of angiogenesis in tumor growth”. eng. In: Seminars in Cancer Biology 3.2 (Apr. 1992), pp. 65–71. ISSN: 1044-579X.

[3] I. P. Tomlinson and W. F. Bodmer. “Modelling the consequences of interactions between tumour cells.” In: British Journal of Cancer 75.2 (1997), pp. 157–160. ISSN: 0007-0920. URL: https://www.ncbi.nlm.nih.gov/pmc/articles/PMC2063276/ (visited on 03/04/2021).

[4] Yuri Mansury and Thomas S. Deisboeck. “The impact of “search precision” in an agent-based tumor model”. eng. In: Journal of Theoretical Biology 224.3 (Oct. 2003), pp. 325–337. ISSN: 0022-5193. DOI: 10.1016/s0022-5193(03)00169-3.

[5] Yuri Mansury, Mark Diggory, and Thomas S. Deisboeck. “Evolutionary game theory in an agent-based brain tumor model: exploring the ‘Genotype-Phenotype’ link”. eng. In: Journal of Theoretical Biology 238.1 (Jan. 2006), pp. 146–156. ISSN: 0022-5193. DOI: 10.1016/j.jtbi.2005.05.027.

[6] Martin A. Nowak. Evolutionary Dynamics: Exploring the Equations of Life. English. Harvard University Press, Sept. 2006.

[7] Deutsch Andreas Basanta David. A Game Theoretical Perspective on the Somatic Evolution of cancer \textbar SpringerLink. 2008. URL: https://link.springer.com/chapter/10.1007/978-0-8176-4713-1_5 (visited on 10/24/2020).

[8] Alexander R. A. Anderson et al. “Microenvironment driven invasion: a multiscale multimodel investigation”. In: Journal of mathematical biology 58.4-5 (Apr. 2009), pp. 579–624. ISSN: 0303-6812. DOI: 10.1007/s00285-008-0210-2. URL: https://www.ncbi.nlm.nih.gov/pmc/articles/PMC5563464/ (visited on 03/20/2020).

[9] David Basanta and Alexander R. A. Anderson. “Exploiting ecological principles to better understand cancer progression and treatment”. In: Interface Focus 3.4 (Aug. 2013). Publisher: Royal Society, p. 20130020. DOI: 10.1098/rsfs.2013.0020. URL: https://royalsocietypublishing.org/doi/full/10.1098/rsfs.2013.0020 (visited on 03/01/2021).

[10] Yiping Hao. Computation and analysis of evolutionary game dynamics. 2013. URL: https://dr.lib.iastate.edu/entities/publication/6429d4ba-6eac-4c83-a8e2-b153acf2a5f6 (visited on 03/07/2025).

[11] Travis I. Zack et al. “Pan-cancer patterns of somatic copy number alteration”. en. In: Nature Genetics 45.10 (Oct. 2013). 00022, pp. 1134–1140. ISSN: 1061-4036. DOI: 10.1038/ng.2760. URL: http://www.nature.com/ng/journal/v45/n10/full/ng.2760.html (visited on 04/03/2014).

[12] Samuel F. Bakhoum et al. “Chromosomal instability drives metastasis through a cytosolic DNA response”. en. In: Nature 553.7689 (Jan. 2018), pp. 467–472. ISSN: 1476-4687. DOI: 10.1038/nature25432. URL: https://www.nature.com/articles/nature25432 (visited on 10/05/2019).

[13] Laurent Sansregret, Bart Vanhaesebroeck, and Charles Swanton. “Determinants and clinical implications of chromosomal instability in cancer”. eng. In: Nature Reviews. Clinical Oncology 15.3 (Mar. 2018), pp. 139–150. ISSN: 1759-4782. DOI: 10.1038/nrclinonc.2017.198.

[14] Kimmel Gregory Ferrall-Fairbanks Meghan. Modeling adaptive therapy in non-muscle invasive bladder cancer \textbar bioRxiv. 2019. URL: https://www.biorxiv.org/content/10.1101/826438v2 (visited on 10/24/2020).

[15] Artem Kaznatcheev et al. “Fibroblasts and alectinib switch the evolutionary games played by non-small cell lung cancer”. en. In: Nature Ecology & Evolution 3.3 (Mar. 2019). Number: 3 Publisher: Nature Publishing Group, pp. 450– 456. ISSN: 2397-334X. DOI: 10.1038/s41559-018-0768-z. URL: https://www.nature.com/articles/s41559-018-0768-z (visited on 03/01/2021).

[16] Ankit Shukla et al. “Chromosome arm aneuploidies shape tumour evolution and drug response”. en. In: Nature Communications 11.1 (Jan. 2020). Number: 1 Publisher: Nature Publishing Group, p. 449. ISSN: 2041-1723. DOI: 10.1038/s41467-020-14286-0. URL: https://www.nature.com/articles/s41467-020-14286-0 (visited on 03/04/2021).

[17] Sohrab Salehi et al. “Clonal fitness inferred from time-series modelling of single-cell cancer genomes”. en. In: Nature 595.7868 (July 2021). Publisher: Nature Publishing Group, pp. 585–590. ISSN: 1476-4687. DOI: 10.1038/s41586-021-03648-3. URL: https://www.nature.com/articles/s41586-021-03648-3 (visited on 03/07/2025).

