## Supplemental figures for "Inverse Game Theory Characterizes Frequency-Dependent Selection Driven by Karyotypic Diversity in Triple Negative Breast Cancer"

### Appendix A

#### *Appendix A.1 ESS convergence across artificially generated datasets*

To assess the velocity of our artificial data towards the ESS, we recorded the distance from the ESS state over time. We generated 1,000 random payoff matrices and solved the replicator equations over time as previously described in Section 2.6. We then verified that the final state was indeed at or near an Evolutionarily Stable Strategy (ESS) by checking the relevant stability conditions. For each solution, we measured the distance of the evolving population state from the ESS at each time point and store these distances. Finally, we compiled all 1,000 distance-to-ESS trajectories into a single figure to visualize how quickly or slowly each population converges toward its ESS.

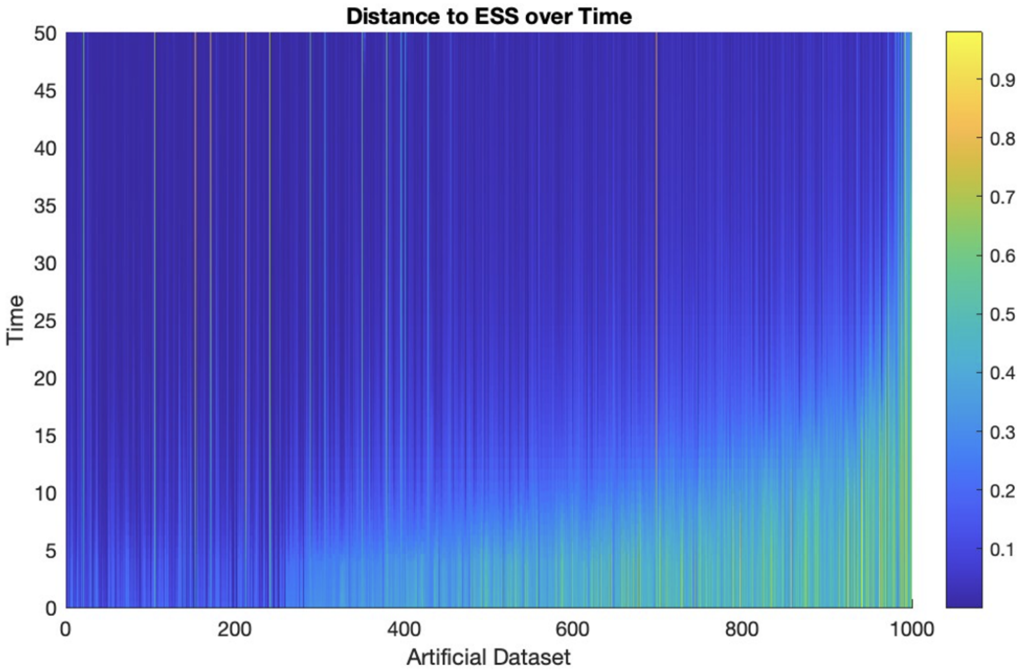


**Figure A1.** Distance to ESS over time for 1,000 artificially generated datasets. The horizontal axis indexes each dataset, while the vertical axis represents time from 0 to 50. Colors denote distance from the ESS (blue indicating smaller distances, yellow larger distances). Each vertical "column" thus shows how quickly and closely a particular solution approaches its ESS over the simulated time span.

#### *Appendix A.2 Inference of zero and non-zero payoff matrix entries via ECO-K in the artificially generated datasets*

We evaluated the ability of the ECO-K framework to infer the underlying matrix structure from a given dataset by randomly generating 1000 payoff matrices as described in Sec. 2.6. For each artificial dataset, we calculated the root mean square error between those entries which were non-zero in either the payoff matrix inferred by our framework $M$ or the underlying matrix $M^*$ (Fig. A2).

We performed this evaluation with varying number of subpopulations and noise levels in the data. Similar to the results obtained from calculating this metric across all all payoff matrix entries (Fig. 2), the recovery algorithm consistently inferred the underlying parametrization of non-zero entries better than would have been expected by random chance (expected $RMSE = \sqrt{\frac{2}{3}}$).

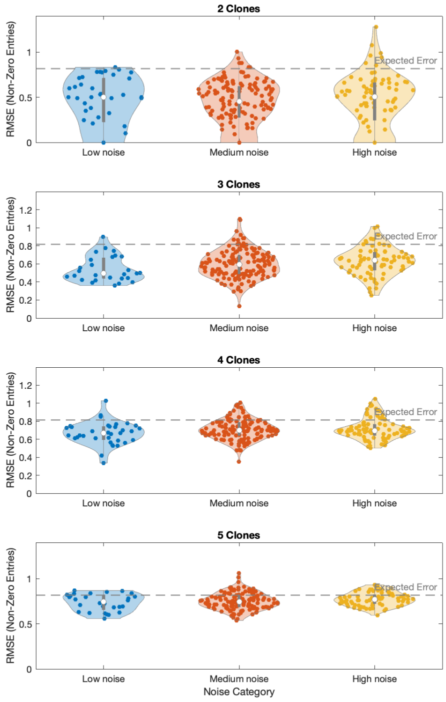


**Figure A2.** Violin plots illustrating the distribution of recovery errors across 1000 artificial datasets, as described in Fig. 2. However, in this plot, we only analyzed entries which were non-zero in either the original payoff matrix, or the matrix which was recovered by ECO-K.

In addition to tracking the overall root-mean-square error (RMSE) between the true and inferred payoff matrices, we also measured how well the algorithm *identifies which entries should be zero*. Concretely, we created two binary masks:

$$\texttt{pm0} = (\texttt{payoff\_matrix} == 0), \quad \texttt{rm0} = (\texttt{recovered\_matrix} == 0),$$

then computed

$$\texttt{all\_sign\_diff}(i) \;=\; \text{RMSE}\Big(\texttt{double}(\texttt{pm0}(:)),\texttt{double}(\texttt{rm0}(:))\Big).$$

In other words, an entry contributed to the error if the original matrix had a zero but the inferred matrix did not (or vice versa). A smaller value for this "inference-of-zeroes" metric indicated that the algorithm more accurately recovered the correct locations of zero versus non-zero entries. By plotting this metric across various noise levels and numbers of SPs (Fig. A3), we evaluated how robustly ECO-K identified which interactions should have remained zero in the presence of measurement noise.

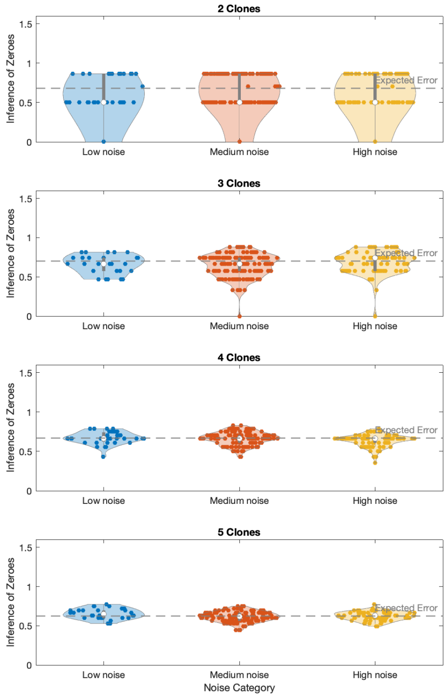


**Figure A3.** Violin plots showing the distribution of the "inference-of-zeroes" metric, computed as

$$\mathrm{RMSE}\Big(\mathrm{double}(\texttt{pm0}(:)),\ \mathrm{double}(\texttt{rm0}(:))\Big),$$

across varying noise levels and numbers of SPs. A smaller value signifies closer agreement between the recovered and true payoff matrices with respect to which entries are zero.
